# Orthorexia is associated with a paradoxical appetitive gastric response to unhealthy foods

**DOI:** 10.1101/2025.06.27.661259

**Authors:** Maya Gumussoy, Evgeniya Anisimova, Serena Lee, Angel Velummylum, Summer Cox, Emily Bagley, Camilla L. Nord, Edwin S. Dalmaijer

**Affiliations:** School of Psychological Science, University of Bristol, Bristol, United Kingdom; Department of Psychology, University of Bath, Bath, United Kingdom; MRC Cognition and Brain Sciences Unit, University of Cambridge, Cambridge, United Kingdom; Department of Psychiatry, University of Cambridge, Cambridge, United Kingdom

**Author notes:** Authors contributed equally (Maya Gumussoy, Evgeniya Anisimova). **Corresponding Author** Dr Edwin S. Dalmaijer, 12a Priory Road, Bristol, BS8 1TU, United Kingdom.

**Keywords:** orthorexia, appetite, disgust, gut-brain interaction, interoception, visceral, somatic, emotion, nausea

## Abstract

Orthorexia involves an obsessive tendency towards “healthy” foods. It is a risk factor for eating disorders (*anorexia nervosa* and the proposed “*orthorexia nervosa*”) and psychological distress, but its biological mechanism remains unknown. We hypothesised that this mechanism operates through the stomach: increased gastric power differentiates appetising from unappetising foods, vagal stimulation reduces food liking, and gastric proto-nausea can be evoked by disgust; all contributors to disordered eating. We used electrogastrography alongside high-density facial landmark tracking to gauge responses to minimally processed (“healthy”), highly processed (“unhealthy”), and culturally unaccepted (“disgusting”) foods in a non-clinical sample (N=77). Trait orthorexia was associated with increased self-reported desire to eat healthy foods. Moreover, we found that higher orthorexia was associated with increasing disgust-related facial responses to unhealthy and disgusting foods. We then identified a relationship between food-healthiness ratings and gastric power: those who generally assigned lower healthiness ratings to foods showed higher gastric power for unhealthy than healthy foods, whereas those who expressed higher healthiness ratings showed higher gastric power for healthy over unhealthy foods. This suggests that the stomach tracks individual differences in eating preferences. Crucially, and contrary to our expectations, higher trait orthorexia was associated with a paradoxical increase in gastric power to unhealthy foods. One explanation is that higher gastric power for unhealthy foods reflects increased appetite for self-denied foods. Alternatively, orthorexia itself could be an adaptive response to exert stronger cognitive control over a pre-existing gastric appetite for unhealthy foods.

## Introduction

Orthorexia is an obsessive tendency towards “pure” foods (Bratman & Knight, 2000; Cena et al., 2019; Fugh-Berman, 2001). It exists as a trait comprised of two highly related factors: an inclination towards healthy foods, and rigid eating behaviours with socio-emotional consequences (Barlow et al., 2024; Barrada & Roncero, 2018). The vast majority of work on orthorexia has been done in cross-sectional questionnaire studies (Meule & Voderholzer, 2021), and work on its underlying biological mechanisms remains notably absent.

In an attempt to empirically study the physiology that contributes to orthorexia, we turned to the stomach. The stomach’s myoelectrical activity increases in power prior to food intake (Levine, 2005), and can differentiate between appetising and unappetising foods (Stern et al., 2001; Zhou & Hu, 2006). On the other side of the spectrum, proto-nausea occurs during episodes of disgust and is primarily characterised by reduced normogastric power (Alladin, Judson, et al., 2024). These associations between stomach rhythm and behaviour could be causal: stimulation of the vagal nerve impacts food liking (Salaris & Azevedo, 2025), and pharmacological normalisation of gastric state reduces disgust avoidance (Nord et al., 2021). Thus, because orthorexia is characterised by appetite for healthy foods and disgust towards unhealthy foods, the stomach is likely involved.

Changes of stomach function and gastric interception occur in many eating disorders (Khalsa et al., 2022; Schalla & Stengel, 2019). Anorexia nervosa is associated with early satiety (Khalsa et al., 2022), whereas bulimia nervosa and binge eating are associated with reduced normogastric power and increased intake before satiation in a water-load task (Van Dyck et al., 2021). However, these are often a consequence rather than a cause of disordered eating (Schalla & Stengel, 2019). Such reversed causality is less likely in orthorexia, which is associated with more selective diet but not necessarily with caloric restriction.

To further reduce the possibility of disordered stomach rhythms, we investigated orthorexic traits in the population rather than in a diagnosed sample. This is also a pragmatic necessity due to the lack of a formal diagnostic category. While the disorder *orthorexia nervosa* has been proposed (Cena et al., 2019; Dunn & Bratman, 2016) and consensus is starting to form on its definition, diagnosis, consequences, and correlated or risk factors (Donini et al., 2022), notable dissent questions the need for a separate diagnosis (Meule & Voderholzer, 2021). Trait orthorexia is associated with trait restrictive eating and a risk factor for disordered restrictive eating (Atchison & Zickgraf, 2022). It is also positively associated with stress, anxiety, and depression symptoms (Barlow et al., 2024), and with disgust sensitivity (Kocabas & Sanlier, 2024). Multi-faceted relationships also exist between disgust and unhealthy foods (Mistry et al., 2025).

In sum, orthorexic traits are distinguishable from other symptoms of disordered eating. We hypothesised that they would be reflected in an appetitive gastric response (increased normogastric power). We also hypothesised orthorexia to be associated with a disgust response towards unhealthy foods, reflected in proto-nausea (reduced normogastric power). We also recorded self-report measures of food perception and disgust, and facial expressions as a behavioural marker of disgust.

## Results

### Self-reported ratings confirm experimental manipulations

We recruited 77 healthy humans (71% women; mean age 25 years, SD=9.7, range=18-63) and presented them with images of foods typically considered “healthy”, “unhealthy”, or “disgusting”. We measured trait orthorexia using the Test of Orthorexia Nervosa (TON-17) questionnaire (Rogowska et al., 2021) [M=43, SD=8.2, range=28-71]. We also asked participants to provide their perceived healthiness ratings for each food stimulus; and ratings of desire to eat, anticipated taste pleasantness of, familiarity with, and disgust towards each food. We followed pre-registered analyses for self-report data to validate our food type classifications (reported in this section) and to test the relationship between desire to eat specific food types and trait orthorexia (see next heading). We conducted linear mixed-effects models with participant and stimulus number accounted for as random effects; using food condition, trait orthorexia, and their interaction as fixed effects.

The “unhealthy” category comprised highly processed foods (e.g. crisps and sausages) for which participants produced lower healthiness ratings compared to the “healthy” category [β=-1.95, *z*=-83.64, *p*=1e-300]. The “disgust” category comprised unusual foods in Western diets (e.g. insects), and also received lower healthiness ratings [β=-1.23, *z*=-52.71, *p*=1e-300], albeit to a lesser extent than “unhealthy” foods [*z*=22.10, *p*=2.84e-108]. “Disgusting” foods were also rated substantially lower on familiarity [β=-1.55, *z*=-59.58, *p*=1e-300] and higher on disgust [β=1.50, *z*=54.82, *p*<1e-300], whereas “unhealthy” foods were no less familiar [β=-0.030, *z*=-1.16, *p*=0.246] or disgusting [β=0.020, *z*=0.72, *p*=0.473] than “healthy” foods.

We also found that participants rated “disgusting” foods as less pleasant than “healthy” foods [β=-1.32, *z*=-43.991, *p*<1e-300] and indicated a lower desire to eat [β=-1.04, *z*=-31.26, *p*=1.53e-214]. “Unhealthy” foods were rated as more pleasant than “healthy” foods [β=0.078, *z*=2.58, *p*=0.010], but did this not translate to a higher desire to eat [β=0.024, *z*=0.73, *p*=0.466].

This confirmed our initial characterisation of foods: “healthy”, “unhealthy”, and “disgusting” food images were indeed rated as such by participants.

**Figure 1.**
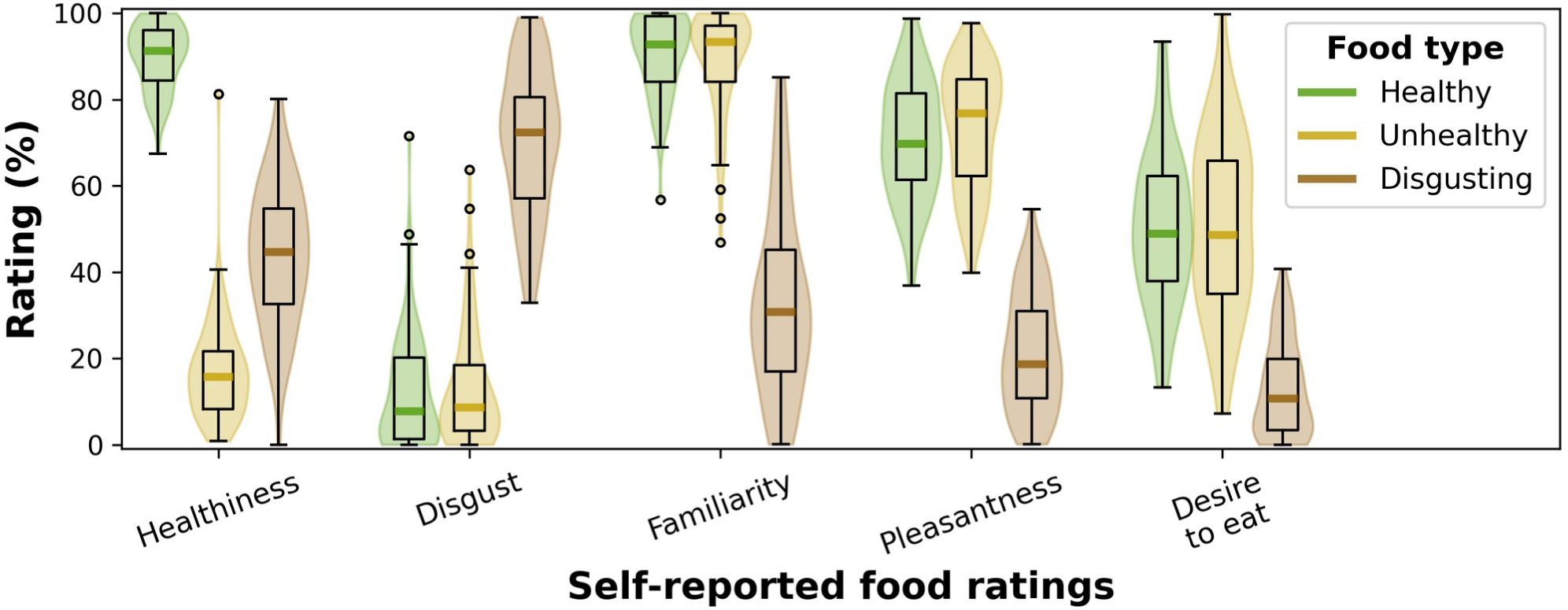
Distributions of ratings offered by participants for foods in three conditions: “healthy” (minimally processed, in green), “unhealthy” (highly processed, in yellow), and “disgust” (stereotypically disgusting in local cultural tradition, in brown). Participants rated 15 stimuli per food condition on healthiness, disgust, familiarity, pleasantness of taste, and desire to eat. Results confirmed the experimental design: “healthy”, “unhealthy”, and “disgusting” food stimuli were indeed perceived as such.

### Trait orthorexia is associated with higher desire to eat healthy foods

Orthorexia did not relate to overall food-healthiness ratings across food types [β=0.029, *z*=0.99, *p*=0.322] or those specific to “unhealthy” stimuli [β=-0.020, *z*=-0.843, *p*=0.399]. An interaction was present for “disgusting” stimuli [β=-0.055, *z*=-2.38, *p*=0.017], such that the difference between healthiness ratings for “healthy” and “disgusting” stimuli was higher for participants with higher orthorexia scores. No statistically significant fixed effects or interactions existed for trait orthorexia and familiarity or disgust ratings.

Higher trait orthorexia was associated with increased overall pleasantness ratings [β=0.118, *z*=3.29, *p*=1.00e-3] and desire to eat [β=0.204, *z*=4.34, *p*=1.45e-5] for all food types. It also interacted with pleasantness [β=-0.100, *z*=-3.31, *p*=9.33e-4] and desire to eat [β=-0.178, *z*=-5.36, *p*=8.46e-8] for “disgusting” stimuli, and with pleasantness [β=-0.138, *z*=-4.60, *p*=4.22e-6] and desire to eat [β=-0.171, *z*=-5.14, *p*=2.75e-7] for “unhealthy” stimuli. That those with higher orthorexia traits reported higher desire to eat all food types was driven by a correlation between orthorexia and desire-to-eat ratings for “healthy” foods [*R*=0.38, *p*=0.001, BF10=31.7], alongside a notable absence of correlation for “unhealthy” foods [*R*=0.05, *p*=0.698, BF10=0.16].

In sum, those with higher trait orthorexia showed a larger difference in healthiness ratings for “healthy” and “disgusting” foods. Trait orthorexia was also associated with larger differences in pleasantness ratings and desire to eat between “healthy” and “unhealthy” foods. This was driven by those higher on orthorexia reporting a higher desire to eat “healthy” foods, but not a reduced desire to eat “unhealthy” foods.

### Higher orthorexia relates to disgust-like facial responses for unhealthy and disgusting foods

In addition to self-reported scores and electrogastrography, we recorded high-density facial landmarks during the presentation of food images, and analysed these with linear mixed-effects models (Figure 2). These again used food type, orthorexia score, and their interactions as fixed effects; and stimulus number as random effect.

**Figure 2.**
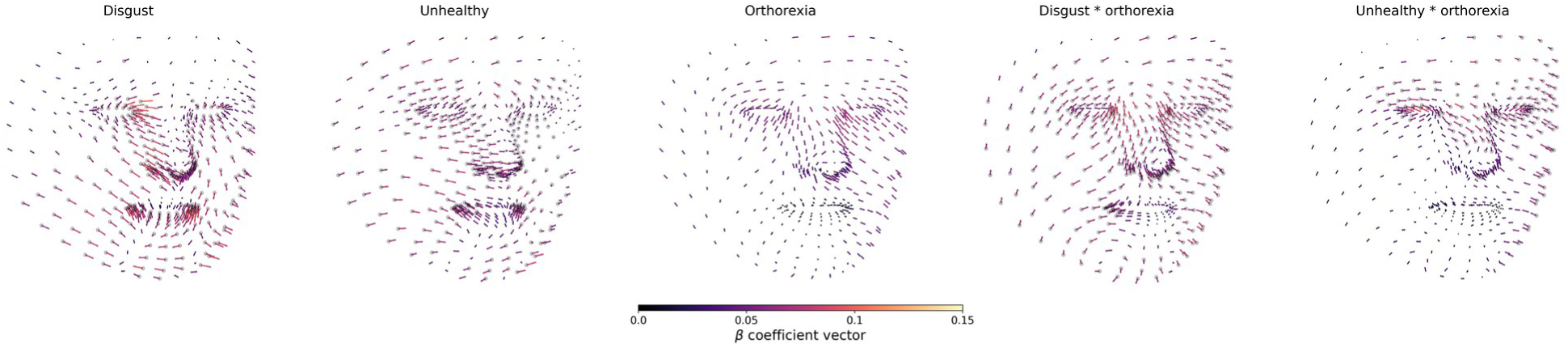
Standardised β coefficient vectors for linear mixed-effects analyses for the relative shift of facial landmarks in three dimensions. Facial responses to “disgust” and “unhealthy” conditions are relative to “healthy” foods, and trait orthorexia was quantified as TON-17 score. Vectors are not plotted to scale of facial movement, but to reflect relative magnitude of coefficients. Facial landmarks are indicated by grey disks, with larger disks indicating statistical significance after Bonferroni correction for multiple comparisons. Findings suggest that individuals higher on orthorexia show more disgust-associated facial expressions to unhealthy and disgusting foods.

Following a pre-registered interest in pairwise differences between food conditions, we observed a partially disgust-associated facial response to “unhealthy” or “disgusting” compared to “healthy” stimuli. After multiple-comparisons correction, there were 287 (out of 468) statistically significant vectors in the “unhealthy” condition and 290 in the “disgusting” condition. Using the Facial Action Coding Scheme (Ekman & Friesen, 1978, 2019), we found facial vectors congruent with action units 1 (inner brow raising), 9 (nose wrinkling), and 15 (lip corner depression) for “unhealthy” foods; and action units 15 (lip corner depression), 16 (lower lip depression), and 25 (lips parting) for “disgusting” stimuli.

Importantly, while no statistically significant vectors were observed for the fixed effect of orthorexia score, we did observe statistically significant vectors for interactions between orthorexia score and food condition: 160 for “unhealthy” and 334 for “disgusting” foods. Relative facial vectors were mainly congruent with action units 4 (brow lowering), 7 (eyelid tightening), 9 (nose wrinkling), 10 (upper lip raising), 41 (eyelid drooping), and 44 (squint). These are highly consistent with the stereotypical facial expression of disgust (Feldman Barrett et al., 2019; Jack et al., 2014; Snoek et al., 2023), suggesting that individuals with higher trait orthorexia showed more disgust-like facial responses to both “unhealthy” and “disgusting” foods.

### Trait orthorexia has an unexpectedly inversed gastric signature

We recorded participants’ stomach rhythms using electrogastrography. Gastric power typically peaks around 3 cycles/minute (0.05 Hz). An increase of this normogastric peak reflects an appetitive response, e.g. in anticipation of food (Levine, 2005). On the other hand, disgust inspires proto-nausea, when gastric power is reduced in the normogastric band and subtly increased in bradygastric (<2 cycles/minute) or tachygastric (4-10 cycles/minute) bands (Alladin, Judson, et al., 2024).

Using a linear mixed-effects model with participant and electrogastrography channel number as random effects, we found reduced gastric power for “unhealthy” [β=-0.033, *z*=-4.07, *p*=4.64e-05] and “disgusting” foods [β=-0.035, *z*=-4.28, *p*=1.87e-05] compared to “healthy” foods. There was also an interaction between frequency and the “disgusting” (compared to the “healthy”) condition [β=-0.017, *z*=-2.16, *p*=0.031], likely reflecting a disgust-specific shift of gastric power from the normogastric to the tachygastric range, i.e. proto-nausea (Alladin, Judson, et al., 2024). Crucially, we also found an interaction between orthorexia scores and the “unhealthy” food condition [β=0.028, *z*=3.54, *p*=4.03e-04] but not the “disgusting” condition [β=-0.001, *z*=-0.08, *p*=940] (both compared to “healthy” stimuli).

This means that, contrary to our expectations, higher trait orthorexia was associated with a reduced difference in gastric amplitude between “healthy” and “unhealthy” foods (Figure 3, top row).

**Figure 3.**
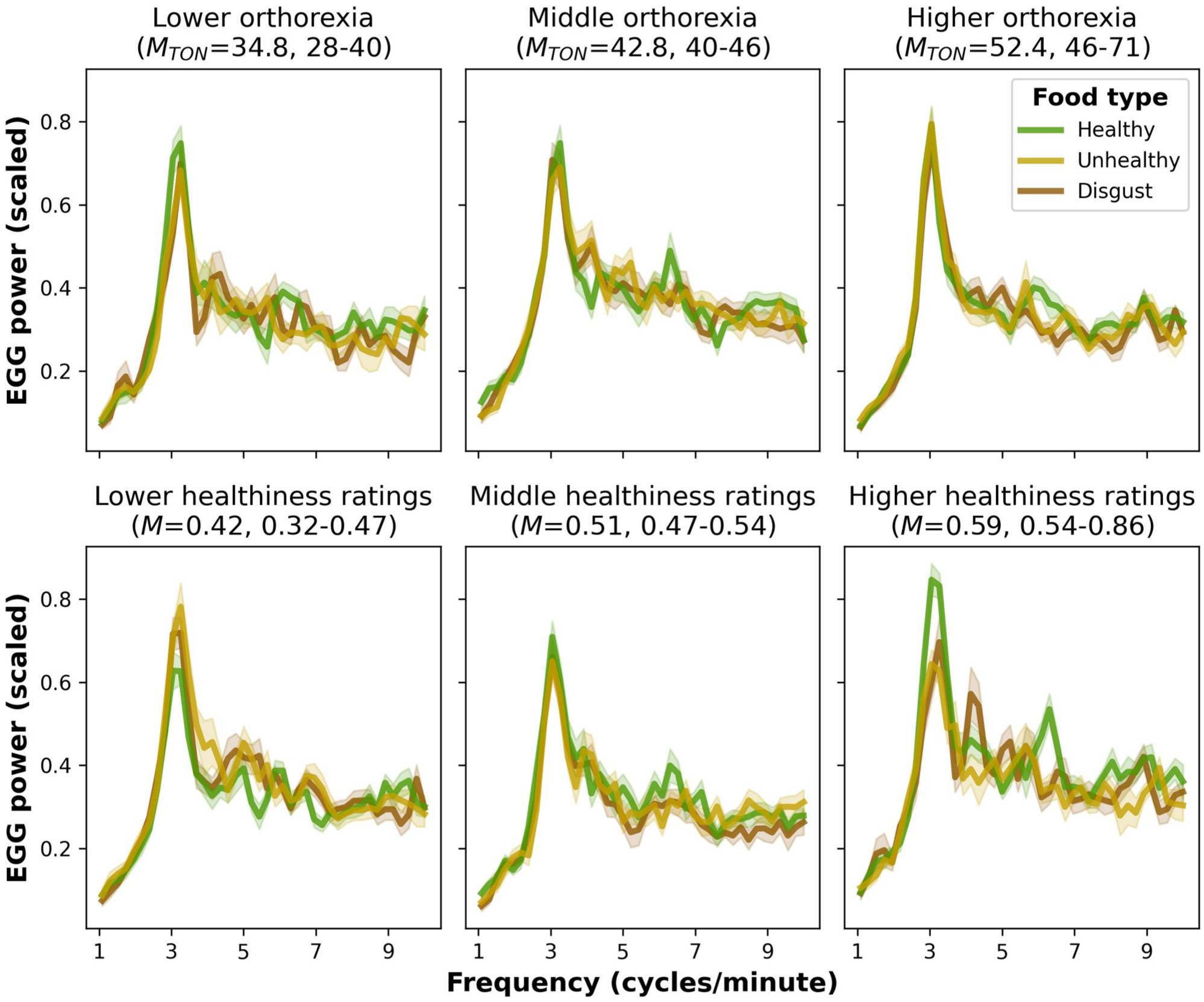
Signal power across the electrogastrographic frequency range during blocks of healthy (green line), unhealthy (yellow line), and disgusting (brown line) food presentation; shown in bins organised by orthorexia (TON-17) score (top row) or by average food-healthiness rating (bottom row). For illustration purposes, power was scaled to individuals’ maximum gastric power in the baseline block. Solid lines reflect the average of participants in each bin (N=26, N=26, N=25), and shaded areas indicate the within-participant standard error of the mean. As is typical for gastric power, peaks are observed at the normogastric frequency of 3 cycles per minute. Counter to expectation, participants with higher trait orthorexia showed lower differences in gastric peak between healthy and unhealthy foods. The ratio in normogastric peak between healthy and unhealthy foods was associated participants’ healthiness ratings for food stimuli, with those who assigned higher healthiness ratings to foods showing higher normogastric power for healthy foods.

### Food-healthiness perception is associated with a congruent gastric response

We expected higher levels of trait orthorexia to inspire an appetitive response to “healthy” foods (increase in normogastric power), and disgust-associated proto-nausea (Alladin, Judson, et al., 2024) to “unhealthy” foods (decrease in normogastric power). Because we found the opposite, we conducted an exploratory analysis to directly test if food healthiness ratings had the expected effect on gastric rhythm. Our approach was similar to the pre-registered linear mixed-effects models (i.e. using participant and electrogastrography channel as random effects), but we replaced orthorexia scores for participant-reported food-healthiness ratings.

We found no difference in gastric signal power between “healthy” foods and “unhealthy” [β=-0.001, *z*=-0.02, *p*=0.987] or “disgusting” foods [β=0.037, *z*=0.95, *p*=0.345]. However, we did find interactions with healthiness ratings for the “unhealthy” [β=-0.102, *z*=-2.89, *p*=3.82e-03] and “disgust” conditions [β=-0.070, *z*=-2.19, *p*=0.028], each compared to the “healthy” condition.

This suggested that higher healthiness ratings were associated with higher ratios of gastric power between “healthy” and “unhealthy” foods. This meant that participants who gave lower food-healthiness ratings across all conditions showed higher gastric power for “unhealthy” than for “healthy” foods. On the other hand, individuals who gave higher food-healthiness ratings showed higher gastric power for “healthy” compared to “unhealthy” foods (Figure 3, bottom row).

## Discussion

Orthorexia is widely recognised as a distinct pattern of food-related thoughts and eating behaviour (Zagaria et al., 2022). Orthorexic traits are associated with psychological distress (Barlow et al., 2024), and may confer risk for restrictive eating disorders including *anorexia nervosa* and the proposed ‘*orthorexia nervosa*’ (Atchison & Zickgraf, 2022). Its biological underpinnings are unknown. Here, we investigated the gastric mechanisms of orthorexic traits. We expected to see it associated with appetitive gastric responses (Levine, 2005; Stern et al., 2001; Zhou & Hu, 2006) (increased normogastric power) to healthy foods, and “proto-nausea” (Alladin, Judson, et al., 2024) (decreased normogastric power) to unhealthy foods. Instead, we found that orthorexic traits were associated with reduced differences in gastric power between healthy and unhealthy foods, suggesting a relative increase in appetitive response to unhealthy foods. This was observed in concert with an association between orthorexia and self-reported desire to eat healthy foods, as well as with heightened facial expressions of disgust for unhealthy and disgusting foods as measured with high-density facial landmark recordings. In short, orthorexia traits were associated with self-reported and facial signatures of disgust, but opposing gastric signatures.

### Orthorexia is associated with facial expressions of disgust

Outside of self-report, few prior behavioural markers of orthorexia have been reported. We conducted facial recordings because we hypothesised that orthorexic traits would inspire disgust to highly processed (“impure”) foods. Restrictive eating disorders have long been associated with increased sensitivity to disgust, especially regarding situations surrounding food and the body (Troop et al., 2002). This is most studied in *anorexia nervosa*, but also holds for orthorexia (Kocabas & Sanlier, 2024). Heightened disgust is theorised to support an increasingly restricted diet as more foods become associated with the emotion of disgust (Anderson et al., 2021). However, much of the evidence supporting this theory relies on self-report and questionnaire data (Aharoni & Hertz, 2012; Hay & Katsikitis, 2014).

The Facial Action Coding Scheme (Ekman & Friesen, 1978, 2019) describes movements of facial muscles as “action units” that combine into facial expressions. While the cultural universality of emotional expressions is a nuanced topic (Feldman Barrett et al., 2019; Jack et al., 2016), the facial expression of disgust has been well characterised in Western populations by its narrowed eyes, wrinkled nose, and parted lips (Darwin, 1872; Jack et al., 2014; Snoek et al., 2023). The action units that most reliably distinguish disgust are 9 (nose wrinkler) and 10 (upper lip raiser) (Snoek et al., 2023), both of which we found associated with orthorexia in response to disgusting and unhealthy foods. We also found stronger narrowing of the eyes in response to disgusting and unhealthy foods as a function of orthorexia. Feelings of disgust, here indexed by facial expressions, could play a key role in driving clinical and behavioural aspects of orthorexia.

### Orthorexia is associated with a paradoxically appetitive gastric signature for unhealthy food

Subjective ratings of food healthiness also supported the general clinical understanding of orthorexia: higher trait orthorexia was associated with higher pleasantness ratings and desire to eat “healthy” foods. Alterations in self-reported responses to food and increased facial expressions of disgust were accompanied by changes in a third dimension: gastric state. We previously demonstrated a causal relationship between gastric state and disgust avoidance (Nord et al., 2021), and hence expected that heightened self-reported and facial expressions of disgust would be accompanied by its gastric signature: proto-nausea, primarily characterised by reduced normogastric power (Alladin, Judson, et al., 2024). However, contrary to our hypothesis, we found orthorexia was associated with an increase in normogastric power (an appetitive response, Levine, 2005) to “unhealthy” foods. In other words, those with high orthorexic traits showed a lower difference in gastric power between “healthy” and “unhealthy” foods.

The paradoxical relationship between orthorexia and gastric power for “unhealthy” foods could represent a surprisingly increased appetitive response to highly processed foods. Perhaps those with stronger orthorexic traits engage in more self-denial of unhealthy foods, which could result in an increased appetitive gastric response towards them. Or the causal direction may run counter: individuals with a pre-existing appetitive gastric response to unhealthy foods could have developed a stricter control of their intake, resulting in stronger orthorexic traits despite enhanced gastric signals. Increased power for “unhealthy” foods might not imply they are more appetitive to those high on orthorexia per se; instead, they might simply fail to differentiate between healthy and unhealthy foods. This would force a more cognitive strategy for food selection, perhaps explaining the association between restrictive eating disorders such as orthorexia and compulsions/fears about losing control of one’s eating (Duradoni et al., 2023; Froreich et al., 2016).

This surprising result may even echo earlier findings in anorexia nervosa, where recovered anorexic patients showed heightened neural reward responses to appetitive rewards (chocolate) (Cowdrey et al., 2011). This was interpreted as increased salience of appetitive food rewards, either contributing to or caused by anorexia nervosa. Others have found a potential association between gastric interoceptive insight and restrained eating in a student sample (Tiemann et al., 2025). Thus, appetitive gastric responses (i.e. increased normogastric power) may represent food ‘salience’ rather than appetite: an increased representation or salience of unhealthy food, which is then interpreted affectively (via facial expression and self-reported) as negatively valenced. These various explanations could be disentangled in future studies, for example by experimentally manipulating gastric power during healthy and unhealthy food viewing and measuring its effects on desirability and facial markers of disgust.

### Limitations

While first introduced as an obsessive tendency towards “pure” and healthy foods (Bratman & Knight, 2000; Cena et al., 2019; Fugh-Berman, 2001), orthorexia is not necessarily unidimensional. Harmful behaviour is captured in the proposed eating disorder “orthorexia nervosa” (Cena et al., 2019), but it has also been argued that “healthy orthorexia” is distinguishable as an eating style (Barrada & Roncero, 2018; Barthels et al., 2019) that is associated with healthy rather than disordered eating (Depa et al., 2019; Roncero et al., 2021; Zickgraf & Barrada, 2021). Indeed, in factor analyses of questionnaires Teruel Orthorexia Scale (TOS) (Barrada & Roncero, 2018) and Test of Orthorexia Nervosa (TON-17) (Rogowska et al., 2021), items probing healthy eating load onto different factors than those probing disorder symptoms. However, despite their supposed separability, orthorexia’s healthy and maladaptive constructs show more overlap than distinction (Anastasiades & Argyrides, 2022) and both are associated with psychological distress (Barlow et al., 2024). Furthermore, reframing healthy eating as “healthy orthorexia” could be seen as unnecessarily medicalising harmless behaviour and risks confusing the proposed diagnostic category of orthorexia nervosa. Nevertheless, our results echo the distinction between healthy eating and orthorexia: self-reported food-healthiness ratings positively correlated with gastric power for healthy over unhealthy foods, whereas orthorexic traits were associated with a reduced difference in gastric power between healthy and unhealthy foods.

At the group level, we observed parting of the lips in response to disgusting foods, but no other disgust-associated action units. This could be due to our choice of stimuli not inspiring strong disgust. For example, we included haggis (in traditional presentation), frog legs, and algae salad; which are contentious but relatively accepted in local diet. Despite this, stimuli in this condition were awarded higher disgust ratings and they induced reductions in normogastric power, another marker of disgust (Alladin, Judson, et al., 2024). This suggests our disgusting stimuli were generally considered as such, but that facial “disgust” responses might be more sensitive to individual differences in food preferences.

Where previous work was able to disentangle anticipatory (visual) and consummatory (flavour) food reward responses (Cowdrey et al., 2011), our visual stimuli were limited to evoking anticipatory responses. While food anticipation is known to increase gastric power (Levine, 2005), it may not necessarily correspond to gastric responses induced by tasting ‘unhealthy’ food categories. Unfortunately, actual food consumption is likely to confound gastric measurements due to the facilitation of normogastric power due to its digestion. Instead, “sham feeding” (holding food in the mouth before spitting it out) (Stern et al., 2001) and visual-only food presentation (Zhou & Hu, 2006) have successfully been used as alternatives.

## Conclusion

In sum, we report behavioural and physiological characterisations of orthorexia: increased facial expression of disgust and increased normogastric power to unhealthy foods. Our findings provide insight into the putative origins of orthorexia: partially related to disgust (via facial expressions), but with distinct (even opposing) features from a typically disgust-associated gastric signature. This provides a first hint of mechanistic insight into the brain-body interactions that drive orthorexic traits. Fuller understanding requires further study, but could ultimately aid development of treatments for pathological symptom levels.

## Methods

### Participants

Our study was approved by the University of Bristol Psychological Science Research Ethics Committee prior to data collection (reference 18057). We recruited a non-clinical sample (N=77) through digital and printed advertisements. Participants provided written informed consent prior to participation, and were reimbursed £10 or with a 1-hour partial course credit after.

Participants were healthy adults (≥ 18 years old) who spoke English, had normal or corrected-to-normal vision, no gastrointestinal issues, and no history of eating disorders. Eighty participants were initially recruited, but three withdrew before the study commenced. The gender distribution skewed towards women (71% women, 26% men, 3% did not disclose), but the relationship between gender and orthorexia is unclear with studies showing no or small gender differences (Barlow et al., 2024; McComb & Mills, 2019; Rogowska et al., 2021; Strahler, 2019; Varga et al., 2013). Self-reported ethnicities were White (58%), Asian (34%), Black (3%), and Mixed (3%); and 3% did not disclose. The sample’s ages ranged from 18 to 63 years, with a mean of 24.9 and standard deviation of 9.71.

### Procedure

Participants were briefed on the experimental procedure and offered room for questions before providing informed consent. Sensors where then placed for electrogastrography (for placement details, see under “*Electrogastrography recording and processing*”). Two researchers were present during electrode application and removal. A web camera was adjusted to centre the participant’s face for facial landmark recording (see under “*Facial landmark recording and processing*”).

The experiment started with a baseline period of ∼3 minutes to stabilise the recording equipment, during which a video was presented (the song “Mahna Mahna” from The Muppets). A passive viewing task followed, in which three blocks of images were presented in randomised order. Each block comprised 30 trials: 2 repetitions of 15 unique images from “healthy”, “unhealthy”, and “disgusting” foods (see under “*Stimuli*”). This block design was necessary to accurately capture the (slow) stomach rhythm. Images were presented for 10s each, preceded by a 500-1000 ms fixation marker, and followed by 500-1000 ms inter-trial interval (blank screen).

After the passive viewing task, participants was asked to rate each stimulus image and to complete the TON-17 questionnaire (see under “*Self*-*report instruments*”). Afterwards, they were debriefed verbally and given a written debrief sheet, including information on local services for wellbeing and eating disorders.

### Stimuli

Stimuli were photos of foods (Table 1) that were categorised as “healthy” if they were unprocessed or minimally processed, or “unhealthy” if they were highly or ultra-processed according to the NOVA classification (Monteiro et al., 2018). While this is a controversial classification system, the foods we selected closely aligned with participants’ perceptions of healthiness (Figure 1). A third type of stimulus was “disgusting” foods, which were uncommon in local culinary tradition.

**Table 1.**
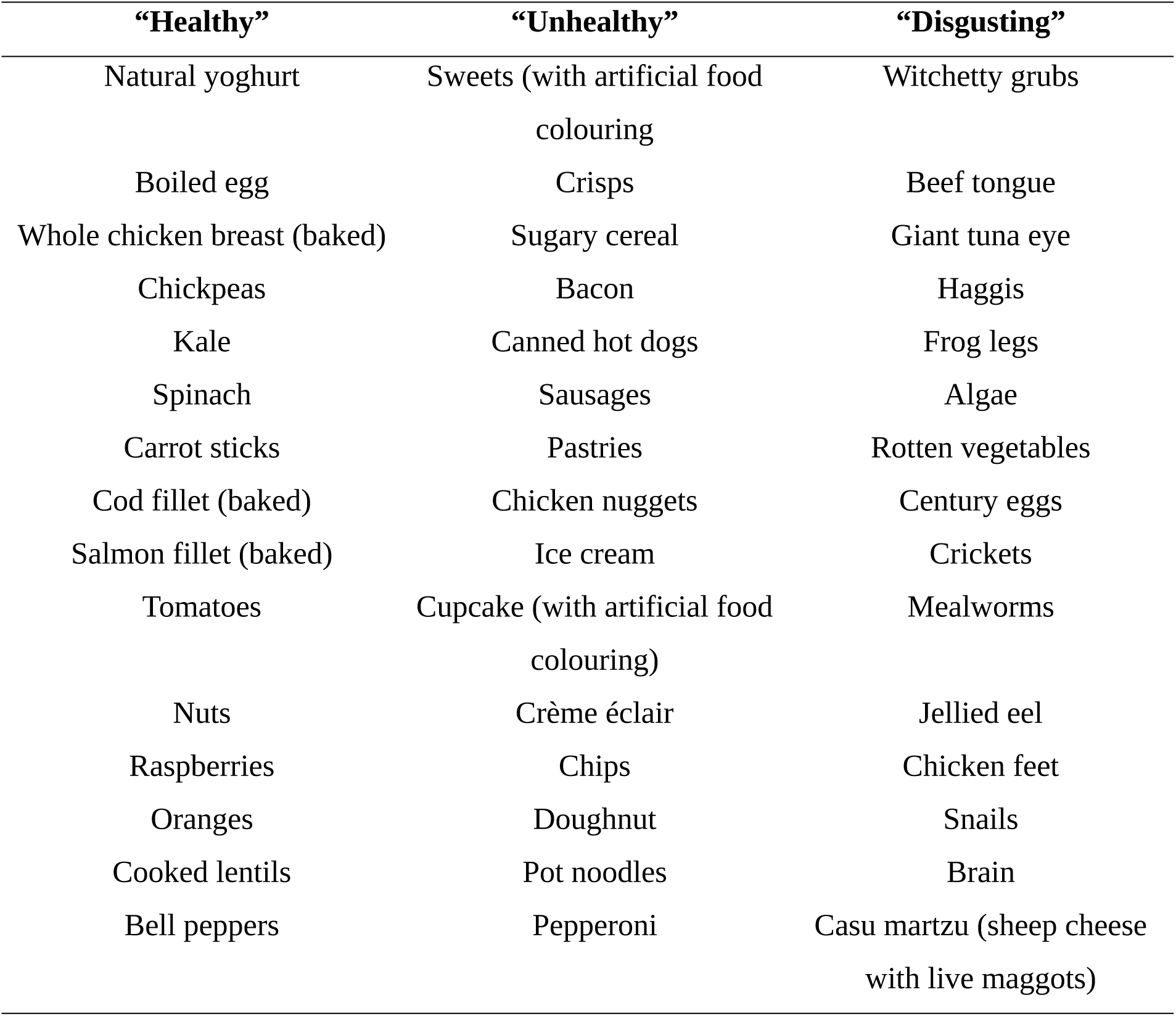
Overview of stimuli for “healthy”, “unhealthy”, and “disgusting” food types.

### Self-report instruments

Orthorexia traits were measured using the 17-item English-language version of the Test of Orthorexia Nervosa (TON-17) (Rogowska et al., 2021), which was developed in accordance with recent recommendations for new orthorexia assessments (e.g., using qualitative data to guide item selection) (Valente et al., 2019) Items are rated on a five-point Likert scale and summed, with higher scores indicating higher orthorexia propensity. Validation of the questionnaire in a Polish sample demonstrated good reliability (Cronbach’s α=0.93) and validity (correlations with questionnaires of orthorexia and eating behaviour) (Rogowska et al., 2021).

At the end of the experiment, participants were asked to rate each stimulus on five different aspects: desire to eat (“How strong is your desire to eat the food in the picture right now?”), disgust (“How disgusted does the thought of eating the food make you feel?”), familiarity (“How familiar are you with the food in the image?”), healthiness (“How healthy do you think the food in the image is?”), and pleasantness (“How pleasant do you expect this food would taste in your mouth?”). The nature and phrasing of these questions follows recommendations in food research (Rogers & Hardman, 2015), and aligns with previous work (Gumussoy et al., 2021; Gumussoy & Rogers, 2023a, 2023b). Responses were captured on a visual-analogue scale in the form of a slider with labelled end-points, and scored between 0 and 1.

### Facial landmark recording and pre-processing

Using OpenCV, we obtained video frames from a webcam with a resolution of 1280×960 pixels at a frequency of 10 Hz. To maintain privacy, video was not recorded but processed immediately. Faces and landmarks were detected using pre-trained convolutional neural networks: BlazeFace short-range (Bazarevsky et al., 2019) followed by MediaPipe FaceMesh (Yan & Grishchenko, 2022). This resulted in the estimation of three-dimensional coordinates for 468 facial landmarks, which were written to file for later analysis.

We preprocessed facial landmarks by setting the origin to the tip of the nose, scaling coordinates to the face width, and aligning each individual face frame with canonical face coordinates with an affine transformation using 7 points on the face (tip of the nose, left and right of the forehead midline, left and right of the chin midline, left and right cheeks). This process ensured that faces were as standardised as possible, to reduce effects of overall rotations over the course of the experiment. We then computed changes in facial landmarks for each stimulus presentation, computing a baseline as mean landmark positions over 3 samples (300 ms) prior to stimulus onset, and differences from that baseline in three directions (horizontal, vertical, and depth) for 13 samples (1.3 seconds, commensurate with duration of emotional expressions (Jack et al., 2014)).

The above resulted in change vectors for 468 facial landmarks over 13 samples, for 3 food type conditions (healthy, unhealthy, disgusting), for 15 stimuli per condition, and 2 presentations of each stimulus. Each of these were fed into linear mixed-effects models (see under heading “Statistical analyses”).

### Electrogastrography recording and pre-processing

Electrogastrography was recorded using a Ganglion amplifier (OpenBCI, NY, USA) with four recording sensors, one reference, and one ground. Sensors were cutaneous Ag-AgCl electrodes (SpesMedica, NeuroTab, DENIS02025), and placed according to established protocol (Al Taee & Al-Jumaily, 2020): the reference just below the xiphoid process; recording sensor 3 halfway between reference and umbilicus, sensor 4 to participant right of sensor 3, sensor 2 in equal measure up and to participant left of sensor 3, and sensor 1 in the same 45° line through sensor 2; and the ground just below the costal margin on the participant’s left side. Channel impedances were checked to be below 10 kΩ before testing.

Participants were seated in reclined position to allow them view of a computer screen while also minimising signal distortion through upright participant position (Jonderko et al., 2005). Before starting the task, participants were briefly shown a trace of their signal to demonstrate the effects of movement, to underscore the instruction to avoid moving and fidgeting.

We recorded data from four sensors, and the following operations were conducted within each channel. To remove offsets (at 0 Hz) in the frequency domain, we mean-centred data. We then computed the signal median and used it to replace any outliers, which were identified as any value over five times the scaled (by 1.4826) median absolute deviation (this is akin to a Hampel filter without a sliding window). To isolate gastric frequencies, we employed a Butterworth filter with a high pass of 0.5 cycles per minute and a low pass of 10 cycles/minute. Finally, a movement filter was applied, which used the target frequency of 3 cycles/minute to estimate and subtract signal attributed to participant movements (Gharibans et al., 2018).

Next, data from all channels was used in an independent component analysis (ICA). This extracted four components, each representing a source contributing to the measured signal. We used a Fournier transform on each component (after applying a Hanning window), and computed signal-to-noise ratio’s as peak normogastric power (within 2-4 cycles/minute) divided by the average power outside of the normogastric domain (0.5-2 and 4-10 cycles/minute). Each component with a signal-to-noise ratio below 3 was zeroed out before the signal was reconstituted in its original space. Six participants showed no component with sufficient normogastric power, and were thus excluded from electrogastrography analyses. The described pipeline closely matches our previous work (Alladin, Berry, et al., 2024).

### Statistical analyses

Our pre-registered analysis plan can be found on the Open Science Framework (https://osf.io/8as4m), and we discuss deviations from it under the next heading. We collected three main types of data: self-report, high-density facial landmark recordings, and electrogastrography.

We took a similar approach across all types of data, using linear mixed-effects models to establish the fixed effects of food type (“unhealthy” and “disgusting”, using “healthy” as reference) and orthorexia (TON-17) score, and their interactions. For electrogastrography data, we also included frequency as fixed effect. We treated participant number as a random effect, and nested random effects for stimulus number (for self-reported ratings and facial landmarks) or sensor number (for electrogastrography).

A benefit of linear mixed-effects analysis is in its statistical power, which depends on both sample size and on the number of trials within participants. In our case, this led to 1155 (self-report), 15015 (facial landmarks), and 11592 (electrogastrography) observations per cell, which aligns with recommendations (Brysbaert & Stevens, 2018).

For facial landmarks, one linear mixed-effects analysis was run per landmark (468) per direction of movement (3). Standardised coefficients for each direction of movement were then combined using Equation 1, and their standard errors were pooled in the same way. Statistical significance was established using Z scores computed via Equation 2. We used Bonferroni correction for multiple comparisons to set our significance threshold to α=0.05/468=1.07e-4.

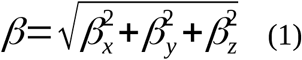

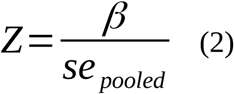

The analysis was coded in Python 3.10.12 (for a tutorial, see Dalmaijer et al., 2025), Brainflow 5.12.0, NumPy 1.26.3 (Harris et al., 2020), SciPy 1.12.0 (Oliphant, 2007; Virtanen et al., 2020), Matplotlib 3.8.2 (Hunter, 2007), and Statsmodels 0.14.1 (Seabold & Perktold, 2010).

### Deviations from the pre-registration

The most obvious deviation from our pre-registration is in sample size. We aimed for N=40, and planned to collect this over two postgraduate dissertation projects (one by authors S.L. and A.V., and another by E.A.). However, due to unexpectedly successful recruitment within each project, we ended up with N=78 (N=77 after exclusion of 1 participant due to recording malfunction).

We made two minor errors in our pre-registration. The first was to include a main effect of “stimulus repetition” for self-reports: while we did repeat stimuli, we only asked for one set of ratings per stimulus. The second error was to list stimulus number (for self-reports), and stimulus repetition and channel number (for electrogastrography) as fixed effects. As their effects on outcomes vary between individuals, these should instead be treated as random effects (Brown, 2021). We now model them as such, nested under participant number.

Following our pre-registered analysis for electrogastrography data, we found that signal power across gastric frequency bands was reduced compared to baseline for “healthy” [β=-0.120, *z*=-14.54, *p*=6.56e-48], “unhealthy” [β=-0.151, *z*=-18.31, *p*=7.55e-75], and “disgusting” foods [β=-0.153, *z*=-18.50, *p*=2.22e-76]. This general reduction was likely an artifact of the experiment’s timeline, in which the baseline always occurred first. Crucially, gastric power was higher for “healthy” foods compared to “unhealthy” foods [*z*=2.74, *p*=6.14e-3], and there was an interaction between the “unhealthy” condition and orthorexia scores [β=0.021, *z*=2.50, *p*=0.013]. The order of other blocks was randomised, so relative differences between conditions could still be meaningfully established. To avoid the baseline artifact, we ran the same analysis without the baseline condition, using the healthy food condition as reference instead. This analysis is reported in the *Results* section, and indeed aligns closely with the findings from the pre-registered analysis listed here.

A more substantial deviation from the pre-registration was the inclusion of an exploratory analysis of the effects of healthiness ratings on electrogastrographic signal. We decided to conduct this after trait orthorexia had an opposite effect on gastric state to what we expected, as it offered a suitable control analysis. Healthiness ratings were a direct measure of individual’s perception of foods’ healthiness, whereas orthorexia was only an indirect measure of this through a general bias towards foods perceived as healthy.

For facial analyses, our pre-registration incorporated pairwise tests of differences between conditions. However, in a multi-conditional design as ours, one should formally consider these as post-hoc tests. We thus decided to run the appropriate multivariate test. This approach aligned with our analysis of self report and gastric measures, and comprised a linear mixed-effects model with participant number as random effect. Fixed effects of condition offered the same information as pairwise comparisons (i.e. “healthy” vs “unhealthy” and “healthy” vs “disgusting” stimulus conditions). Further, the updated approach allowed us to include TON-17 scores to quantify the fixed effect and interactions of trait orthorexia. We aligned with the pre-registration in running analyses per plane of movement (horizontal, vertical, and depth) before combining them. We also followed our pre-registered Bonferroni correction for multiple comparisons.

### Open materials and data

We deposited anonymised data on Zenodo (https://doi.org/10.5281/zenodo.15722062), from where in can be accessed publicly. We made our analysis code freely available through GitHub (https://github.com/esdalmaijer/2025_orthorexia_manuscript), and we also archived a version on Zenodo (https://doi.org/10.5281/zenodo.15722842) for persistence.

## Acknowledgements

At the time of testing, author EA was enrolled in the *MSci in Psychology and Neuroscience* and authors SL and AV in the *MSc in Applied Neuropsychology* at the University of Bristol. CLN and EB are supported by the Medical Research Council (MC_UU_00030/12) and a Wellcome Career Development Award (226490/Z/22/Z).

## Author contributions

Conceptualisation: MG, ESD; Data curation: ESD; Investigation: EA, SL, AV, SC; Methodology: MG; EA, ESD; Software: ESD; Supervision: MG, ESD; Visualisation: EA, ESD; Writing (original draft): MG, EA, ESD; Writing (review and editing): MG, EA, SL, AV, SC, EB, CLN, ESD.

